# chewBBACA: A complete suite for gene-by-gene schema creation and strain identification

**DOI:** 10.1101/173146

**Authors:** Mickael Silva, Miguel Machado, Diogo N. Silva, Mirko Rossi, Jacob Moran-Gilad, Sergio Santos, Mario Ramirez, João André Carriço

## Abstract

Gene-by-gene approaches are becoming increasingly popular in bacterial genomic epidemiology and outbreak detection. However, there is a lack of open-source scalable software for schema definition and allele calling for these methodologies. The chewBBACA suite was designed to assist users in the creation and evaluation of novel whole-genome or core-genome gene-by-gene typing schemas and subsequent allele calling in bacterial strains of interest. The software can run in a laptop or in high performance clusters making it useful for both small laboratories and large reference centers. ChewBBACA is available at https://github.com/B-UMMI/chewBBACA or as a docker image at https://hub.docker.com/r/ummidock/chewbbaca/.

**DATA SUMMARY:**

1. Assembled genomes used for the tutorial were downloaded from NCBI in August 2016 by selecting those submitted as *Streptococcus agalactiae* taxon or sub-taxa. All the assemblies have been deposited as a zip file in FigShare (https://figshare.com/s/9cbe1d422805db54cd52), where a file with the original ftp link for each NCBI directory is also available.
2. Code for the chewBBACA suite is available at https://github.com/B-UMMI/chewBBACA while the tutorial example is found at https://github.com/B-UMMI/chewBBACA_tutorial.

**I/We confirm all supporting data, code and protocols have been provided within the article or through supplementary data files. ⊠**

**IMPACT STATEMENT:** The chewBBACA software offers a computational solution for the creation, evaluation and use of whole genome (wg) and core genome (cg) multilocus sequence typing (MLST) schemas. It allows researchers to develop wg/cgMLST schemes for any bacterial species from a set of genomes of interest. The alleles identified by chewBBACA correspond to potential coding sequences, possibly offering insights into the correspondence between the genetic variability identified and phenotypic variability. The software performs allele calling in a matter of seconds to minutes per strain in a laptop but is easily scalable for the analysis of large datasets of hundreds of thousands of strains using multiprocessing options. The chewBBACA software thus provides an efficient and freely available open source solution for gene-by-gene methods. Moreover, the ability to perform these tasks locally is desirable when the submission of raw data to a central repository or web services is hindered by data protection policies or ethical or legal concerns.

## INTRODUCTION

Read mapping approaches using Single Nucleotide Polymorphisms (SNP)/Single Nucleotide Variants (SNV) have been widely used for studying bacterial genomes [1]. However, gene-by-gene (GbG) approaches have also been advocated in the context of genomic epidemiology as an expansion of Multilocus Sequence Typing (MLST) [2] allowing portability, scalability, and independence from a defined reference strain. For these reasons, GbG increasingly gains popularity and has been adopted by PulseNet International as the method for bacterial strain discrimination using high throughput sequencing [3]. GbG relies on comparing the draft genome of a strain of interest against a pre-defined schema, typically using a BLAST [4] based approach. This schema can be composed of core loci, which are present in all or the great majority (e.g. 95%) of the analyzed strains (core genome MLST schemas or cgMLST), or including all loci detected in the strains of interest. The latter are referred to as whole genome or pan genome MLST schemas (wgMLST or pgMLST).

A locus in a schema can be a complete coding sequence (CDS) or a subsequence of it, as in traditional MLST. Defining a locus as a CDS, allows linking the variability found to potential changes in proteins and thus, with phenotype. The definition of locus is currently dependent on the algorithm used for comparing loci and defining the schema, hampering the comparison between different GbG approaches.

Only few software are available for GbG allele calling and no tools are available for schema creation and validation. Two commercial platforms offer GbG analyses: Ridom SeqSphere+ (http://ridom.de/seqsphere/) and Bionumerics (http://www.applied-maths.com/applications/wgmlst). Since these are proprietary, closed source software, their GbG allele calling algorithms are incompletely described [5–6], although Ridom schemas have been publicly released (http://www.cgmlst.org/).

BIGSdb was the first open-source freely available platform allowing cgMLST analysis [7] and is currently the basis of the PubMLST website (https://pubmlst.org/). More recently, EnteroBase provides comprehensive cgMLST and wgMLST schemas and an allele calling engine for three major foodborne bacterial pathogens (https://enterobase.warwick.ac.uk/). A limitation of EnteroBase is the requirement to submit reads to the website or to public repositories (NCBI SRA/EBI ENA), since currently no stand-alone versions of their allele calling algorithm are available. At present, the only published open-source stand-alone GbG allele calling algorithm is Genome Profiler [8] which, however, uses a single CPU core making it unsuitable for large scale analyses.

We developed chewBBACA to be a complete stand-alone pipeline for GbG analyses, including constructing and validating novel cg/wgMLST schemas and performing CDS allele calling suitable for large scale studies.

## THEORY AND IMPLEMENTATION

chewBBACA is composed of three main modules: Schema Creation, Allele Calling and Schema Evaluation. These modules can be used together in order to define and evaluate new wg/cgMLST schemas for species of interest. A general workflow of such process is presented in Fig. 1.

**Figure 1.**
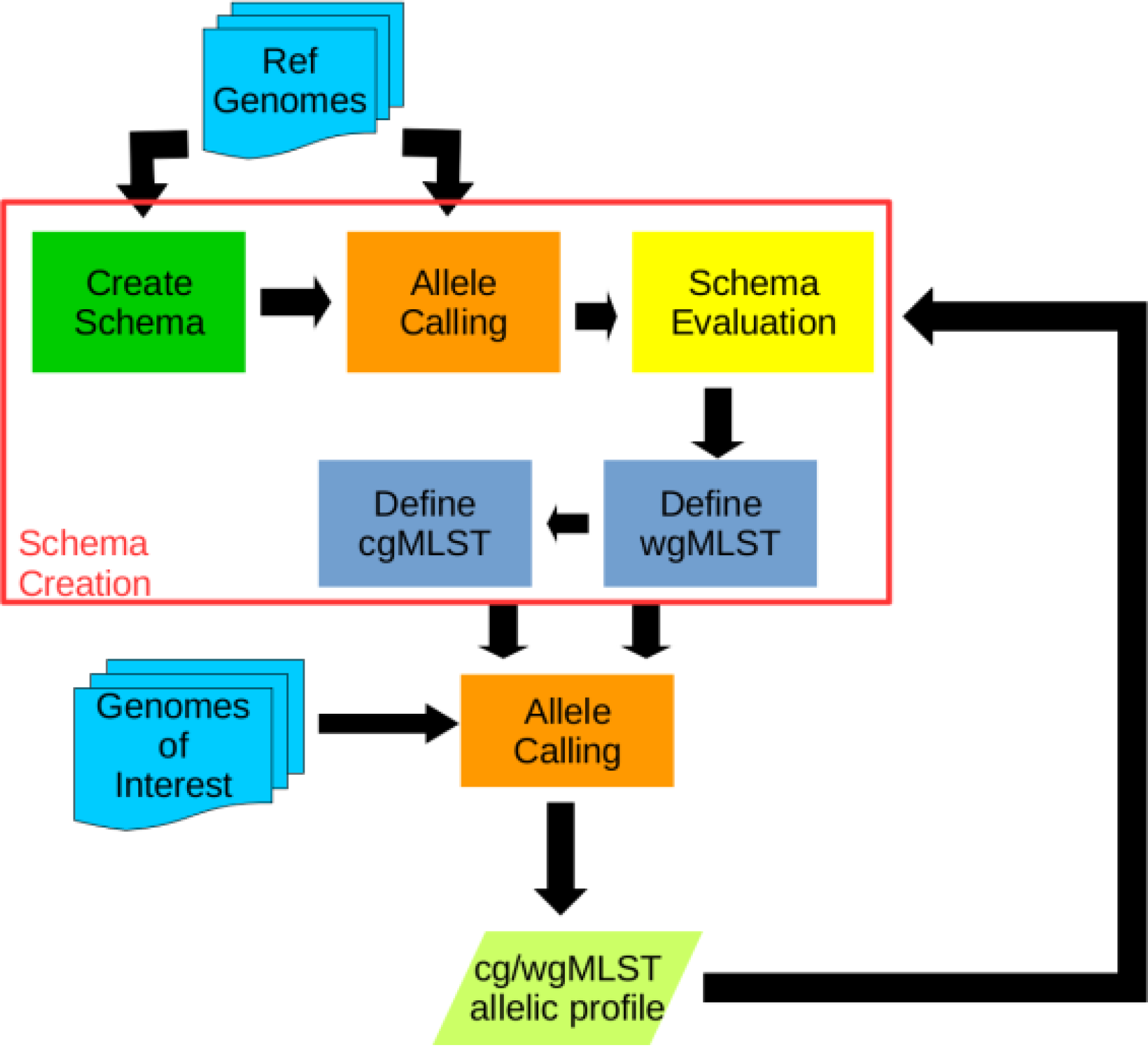
chewBBACA workflow from schema definition to schema evaluation

### wg/cgMLST Schema creation

The First module is the Schema Creation, which allows the definition of wg/cgMLST schemas from user provided complete genomes or draft assemblies, focusing on excluding paralogous loci, detection of contaminated/poor quality assemblies and supporting user decisions towards the identification of the most appropriate schema through interactive graphic data analysis. This module uses an iterative approach for CDS comparison in the selection of loci that is more computationally efficient than the Markov Clustering Step typically used in software such as OrthoMCL [9] or CD-hit [10]. In order to create a wgMLST schema, the user provides a set of genomes in FASTA format. The algorithm first defines the CDSs of each genome using Prodigal [11]. In the next step, all the CDSs in the genomes are compared in a pairwise fashion, resulting in a single file containing all unique CDSs identified in the genomes. This comparison is a two-step process. Firstly, all the CDSs having identical sequence of other CDSs but being smaller in length are removed and the larger CDS is kept. At the same time, the algorithm also removes all CDSs with a length less than indicated in the “-l” parameter. In the second step, the remaining CDSs are clustered in unique loci by performing an all-against-all BLASTP search and calculating the Blast Score Ratio (BSR) [12]. CDSs with a BSR pairwise comparison equal or greater than 0.6 are considered alleles of the same locus and the larger allele (in bp) is kept in the list. This procedure defines the schema as a set of CDSs, each representing the largest single allele of distinct loci. The Allele Calling module is then used to populate the schema with alleles using the same genomes used for its creation. This step allows the identification and exclusion of possibly paralogous loci. The Allele Calling algorithm detects if a CDS in the genome under analysis matches more than one locus in the schema, indicating that those loci can be paralogous. The Allele Calling module outputs a list of such loci to be removed from the wgMLST schema or to be further investigated. From the created wgMLST schema, cgMLST schemas can be defined by selecting the loci that are present in a predetermined percentage of the analysed strains, typically 95%-99%.

### Allele calling Algorithm

The Allele Calling algorithm is based on CDSs identified by Prodigal [11] with similarity determined using a BLASTP BSR approach, allowing the detection of alleles with divergent DNA sequences but similar encoded proteins. This allows the identification of alleles that would be considered absent loci with BLASTN, while retaining the full diversity found at the DNA sequence level. The algorithm is defined as presented in Fig. 2. A BLASTP database is created, containing all the translated CDSs identified by Prodigal in the query genome. A 100% DNA identity comparison is performed on all the genome of interest CDSs against each locus allele database. If an exact match is found, an allele identification is attributed to the CDS (and tagged as *EXC – Exact Match*, in the statistics output). If not, a BLAST BSR approach is used to identify the allele. To improve computational efficiency, chewBBACA performs the similarity search on each locus in the schema separately, parallelising the jobs using the specified number of CPUs. For each locus, a short list containing the most divergent alleles is queried against the BLASTP database. The BSR is calculated for each hit and based on these results and a size validation step, the locus is either considered not found (tagged as *LNF – Locus Not Found*) or a new allele of the locus is inferred. The size validation step excludes alleles larger than or smaller than 20% of the locus allele length mode (Defined as *ASM – Alleles Smaller than Mode* or *ALM – Alleles Larger than Mode*) (Fig.3a). Furthermore, the identification of loci as duplicated in the genome of interest is also reported. Such matches are identified as *Non-Informative Paralogous Hits* (*NIPH*), if at least two CDSs have best matches with alleles of the same locus but presenting less than 100% identity, or *NIPHEM – NIPH Exact Match* if 100% identity to existing alleles is detected (Fig. 3b). Futhermore, the algorithm detects whether the CDS match is close to the 5’ or 3’ ends of a contig and a larger allele that contains the matched sequence would exceed the contig length. Such sequences are tagged as *Possible Locus On the Tip* (*PLOT*) (Fig. 3c). Finally, the Allele Calling module identifies possible paralogous (as described above) checking if there are CDS matching alleles in two or more different loci (Fig. 3d). After running each genome, the loci database is updated with the newly found alleles and, whenever required, a locus short-list is also updated with a new divergent allele.

**Figure 2.**
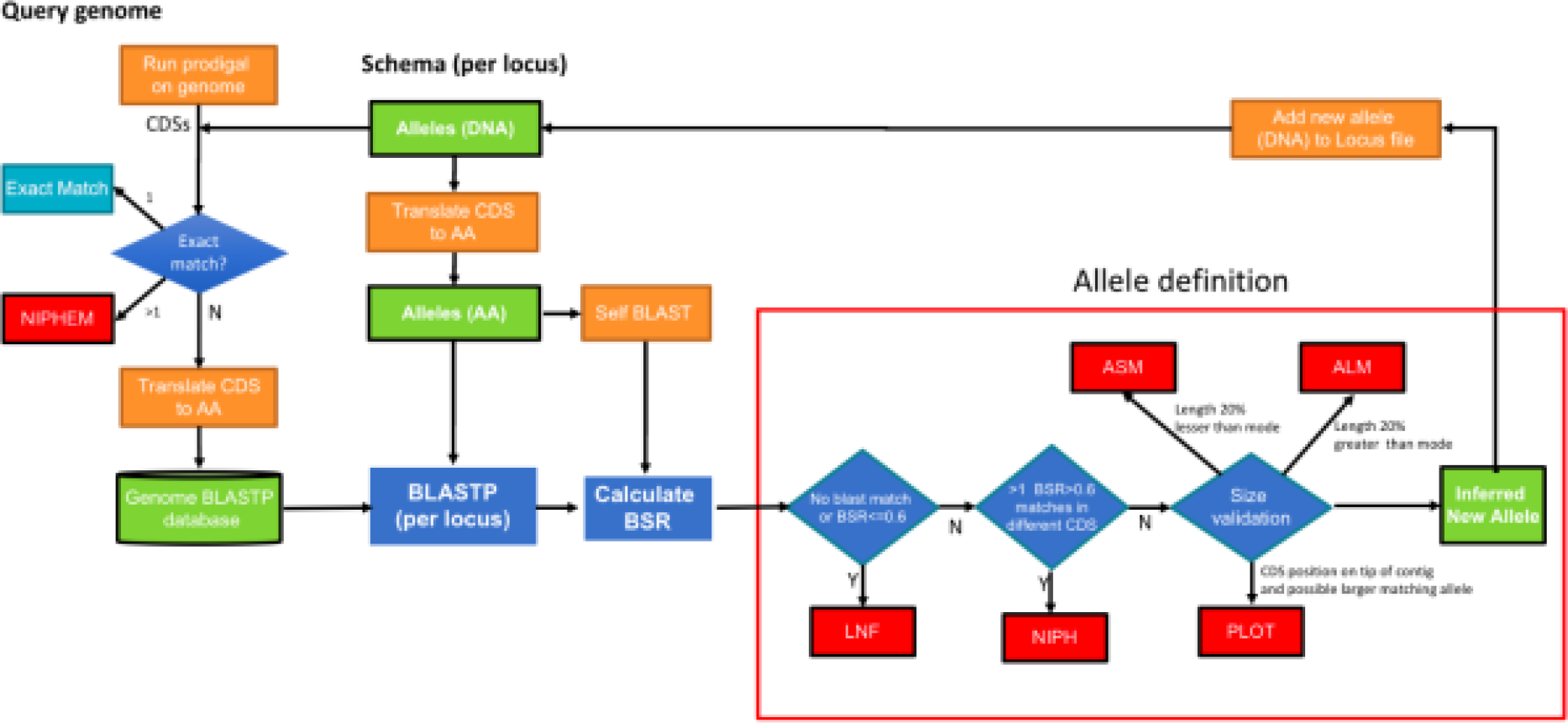
chewBBACA allele callign algorithm.

**Figure 3.**
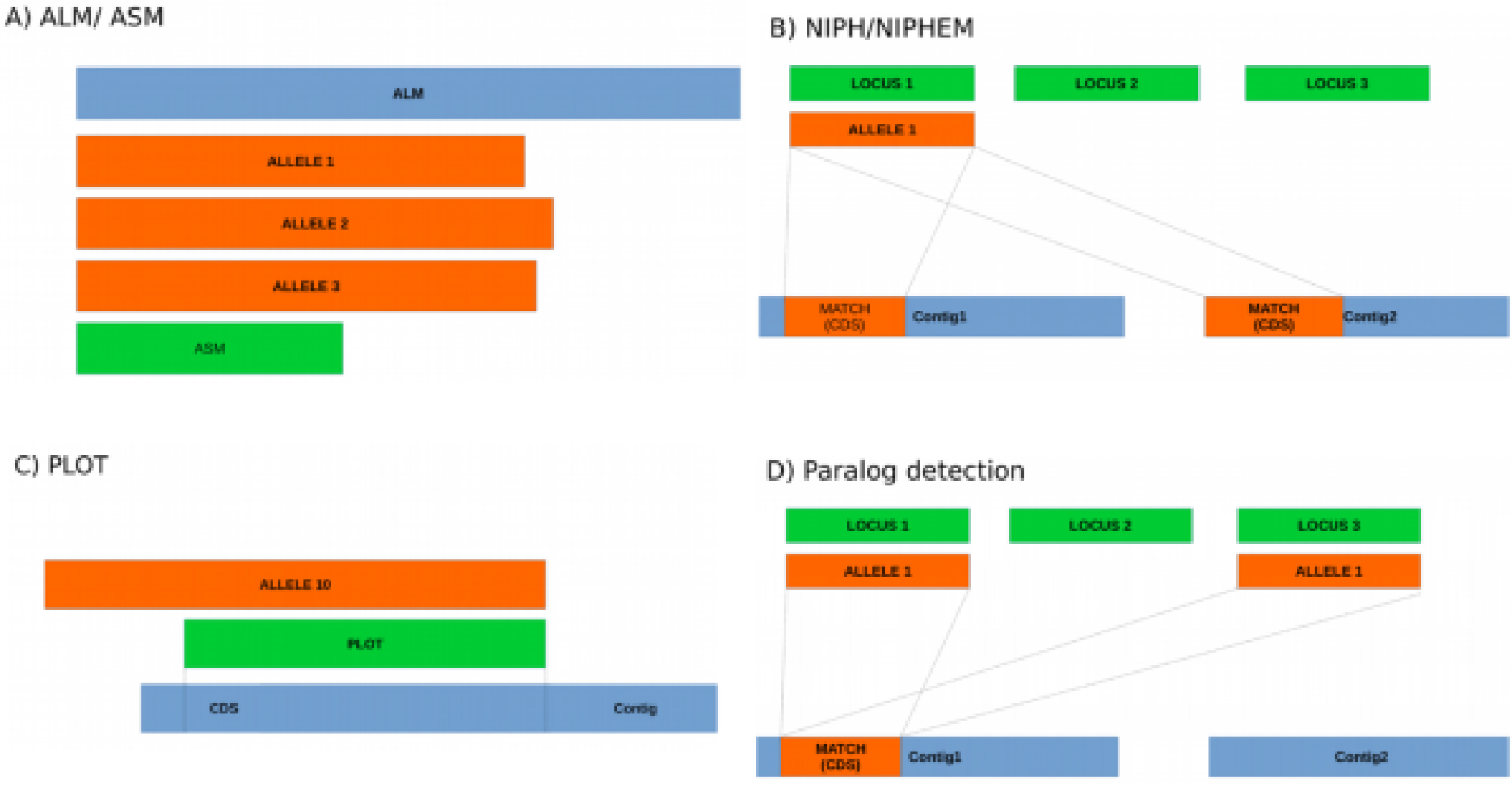
chewBBACA Allele definition outputs. A) size exclusion of alleles 20 % smaller or larger than the allele length mode for the loci B) Detection of loci duplication on the draft genome C) detection of locus identified on the 5’ or 3’ ends of the contig D) Detection of paralogous loci

### Schema Evaluation

This module allows the assessment of the suitability of including each locus in a schema through a suite of functions to graphically explore and evaluate the type and extent of allelic variation detected in each of the chosen loci. This module also creates multiple sequence alignments of the alleles of each locus using MAFFT [13] and constructs neighbour-joining trees using ClustalW2 [14], allowing the exploration of the potential consequences of the variability at each locus. This module can be used to analyse any existing cg/wgMLST schema, including those created by other methodologies, since the analysis input is a set of FASTA files, one per locus, with all identified alleles.

A more complete description of each module and their functionalities is available at https://github.com/B-UMMI/chewBBACA/wiki

### Benchmark

The performance of chewBBACA’s allele calling algorithm was evaluated for *Streptococcus agalactiae* assemblies (~2Mb genome) using a cgMLST schema of 1,264 loci. Benchmarks were performed on a high-performance cluster (HPC) with Intel^®^ Xeon™ E5-2630 v4 @ 2.20GHz CPUs, up to 256Gb RAM and an SSD distributed storage in RAID1; a laptop with Intel^®^ Core™ i5-7200U @ 2.50GHz × 4 CPUs, 8Gb RAM and a NVMe SSD storage; and a laptop with Intel^®^ Core™ i7-3630QM @ 2.40GHz × 8 CPUs, 8Gb RAM and a SATA2 HDD storage. Allele calling was conducted for 100 *S. agalactiae* assemblies in the HPC cluster using 2 to 40 CPUs, and for a subset of 50 assemblies in both laptops using 2 and 4 CPUs (Fig 4). Each CPU data point was run 5 times. In terms of CPU performance, the time it took to run each sample decreased almost linearly up to 16 CPUs, at which point disk possibly I/O storage access becomes the bottleneck and no increase in performance is observed. At peak performance, allele calling takes approximately 4 seconds for each sample. Since this algorithm is I/O intensive, a substantial increase in performance can be observed in a laptop with SSD storage, in comparison to another with HDD storage. In any case, allele calling in a laptop using 4 CPUs can be performed for each sample in approximately 9 seconds with SSD storage and 15 seconds with HDD storage.

**Figure 4.**
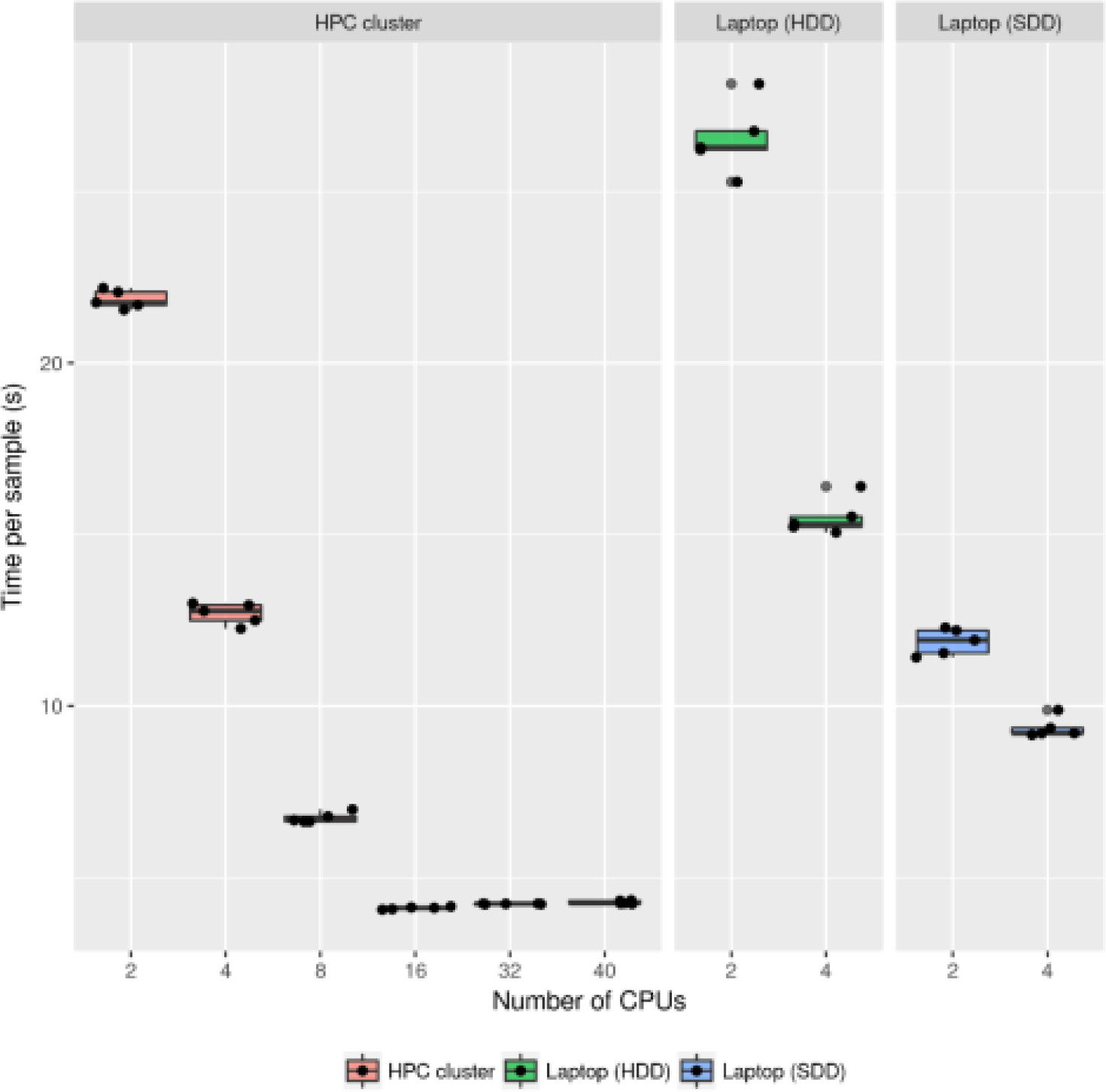
Benchmarking of chewBBACA’s allele calling algorithm for bacterial genome assemblies (^~^2Mb) using a cgMLST schema of 1264 loci on a HPC cluster and two laptops with different storage devices. The allele calling was executed 5 times for each CPU data point.

### Usage Example

A tutorial providing a complete usage example, demonstrating the creation of a schema for *Streptococcus agalactiae*, from publicly available complete genomes and assemblies available at NCBI/ENA is provided at https://github.com/B-UMMI/chewBBACA_tutorial.

## CONCLUSION

The chewBBACA suite was developed to allow performing GbG analyses in high-end Unix based laptops but chewBBACA can also be easily run in HPC, facilitating its adoption into large-scale automated analysis pipelines. The good performance of the software in current laptops and in HPCs, allows a flexible implementation in small laboratories or large reference centres. The allele calling engine of chewBBACA uses FASTA files with draft assemblies or complete genomes as input and returns as output an allelic profile matrix and a set of FASTA files containing the full allelic diversity of each locus. Currently available cg/wgMLST schemas, can be adapted to run using chewBBACA’s allele calling engine. The chewBBACA suite is the first to provide schema creation tools and to enforce CDS allele calling, which can be important to evaluate phenotypic diversity including the identification of the potential mechanisms underlying the success of particular clones. Since there is an urgent need for bioinformatics solutions that will allow the development of nomenclature-based schemas [15], future work will focus on centralised repositories for schemas and allele definitions that can be synchronised with local allele calling outputs to facilitate the development of common schemas and nomenclatures for cg/wgMLST allowing a more widespread application of GbG methodologies in public health.

## AUTHOR STATEMENTS

### Acknowledgements

The authors would like to thank Eduardo Taboada and Peter Kruczkiewicz from National Microbiology Laboratory at Lethbridge, Public Health Agency of Canada, Lethbridge, Alberta, Canada., for insightful discussions.

## ABBREVIATIONS

SNP, Single Nucleotide Polymorphisms; SNV, Single Nucleotide Variants; GbG, gene-by-gene; cgMLST, core genome MLST; wgMLST, whole genome MLST; pgMLST, pan genome MLST; CDS, coding sequence; BSR, Blast Score Ratio;

